# Ancient DNA Clarifies the Identity and Geographic Origin of the holotype of the genus *Ctenomys*

**DOI:** 10.1101/2024.09.10.612281

**Authors:** Renan Maestri, Gislene Lopes Gonçalves, Violaine Nicolas-Colin, Anna Bryjova, Rodrigo Fornel, Eric Coissac, Pierre Taberlet, Gilson Rudinei Pires Moreira, Thales Renato Ochotorena de Freitas

## Abstract

*Ctenomys* Blainville 1826 ranks among the top ten most diverse mammal genera in terms of species richness. However, the taxonomic history of *Ctenomys brasiliensis* Blainville,1826, the corresponding type species, has long been obscured by a dearth of information regarding the collection data of the type material, compounded by an elusive geographic origin. Here, employing ancient DNA methodology, we sequenced the complete mitogenome of the remaining type specimen and conducted an extensive historical investigation to correlate originally described locality names with present-day locales in South America. Our analysis unequivocally confirms that the type specimen corresponds to the species currently designated as *Ctenomys minutus* Nehring, 1887. This resolution lays to rest a century-old debate surrounding the provenance of the type specimen, rejecting prior hypotheses that placed its collection site in southeastern Brazil or Uruguay. Instead, our evidence suggests it was likely obtained from a third location in southernmost Brazil. Previous analyses overlooked this new location due to confusion surrounding geographic nomenclature and labeling errors, issues rectified by our combined mitogenomic and historical approach. Furthermore, quantitative morphological analyses boost our findings, demonstrating a closer affinity between *C. brasiliensis* and *C. minutus* within the same species group. Accordingly, we validate *C. brasiliensis* and propose *C. minutus* as its junior synonym. Our study underscores the importance of robust DNA analyses in confirming the identity and geographic origins of type specimens, especially for *Ctenomys* species with similar phenotypes, and specimens collected centuries ago.

## INTRODUCTION

Identifying the geographic location of a type specimen poses significant challenges, particularly for ancient species descriptions characterized by doubtful or incomplete collection data. This challenge commonly arises due to the absence of available geographic coordinates, and the continual evolution of location names over time, often resulting in broadly defined descriptors such as municipalities, states/provinces, or even countries. Additionally, the lack of DNA or difficulties in extracting DNA for old preserved specimens presents further obstacles to accurately identifying the type species and determining its evolutionary relationship to subsequently described taxa. In some cases, these challenges may prove insurmountable, contingent upon the level of information available in museum records and species descriptions, as well as the preservation method employed in scientific collections (e.g., taxidermy, alcohol, formalin, etc.), which may vary depending on the particular characteristics of the animal or plant (McGuire et al. 2018). Furthermore, older preserved material often presents greater difficulty in DNA extraction from remaining tissues and may feature outdated, incomplete, or non-standardized species descriptions.

Mammals preserved decades ago were typically taxidermized, with their skins treated using borax or a similar substance, while their bones, primarily the cranium and mandible, were commonly preserved as such. The scarcity of fresh material poses a significant challenge in extracting DNA from ancient mammals housed in museums and scientific collections. Nonetheless, DNA analyses of museum specimens have proven to be a valuable resource for discovering new species and genera (Castañeda-Rico et al. 2022), as well as yielding numerous insights into evolutionary and systematics relationships (de Abreu-Jr et al. 2020; Yuan et al. 2022). However, extracting DNA from holotypes presents its own set of challenges, as it involves balancing the need to safeguard invaluable specimens against the potential damage caused by extracting DNA samples (Castañeda-Rico et al. 2022). Nevertheless, DNA extraction from holotypes has emerged as a crucial source of information for refining the taxonomy and its systematic relationships within a group (Pagès et al. 2010; Nations et al. 2022; Ruelas and Pacheco 2022).

The genus *Ctenomys*, recognized as the most species-rich among subterranean mammals and listed among the top ten most diverse mammal genera (Mammal Diversity Database 2024), was proposed in 1826 by Blainville, who designated its type species as *Ctenomys brasiliensis*. Since then, over 65 additional species have been described (D’Elía et al. 2021; Brook et al. 2022; Mapelli et al. 2023; Sánchez et al. 2023; Teta et al. 2023; Verzi et al. 2023; Alvarado-Larios et al. 2024; Mammal Diversity Database 2024; Tammone 2024). The distribution of the entire genus spans the southern half of South America (Freitas et al. 2021). Phylogenetic and paleontological analyses suggest this remarkable diversity emerged within a mere 5 million years (Upham and Patterson 2015; Verzi et al. 2021), constituting one of the most rapid mammalian radiations on record.

Despite centuries of research spanning various fields of knowledge (Freitas et al. 2021), the origin of the first collected specimens of *Ctenomys* remains elusive. Blainville (1826) described the holotype in the French publication “Annales des Sciences Naturelles” published by the National Museum of Natural History of Paris (MNHN). In the publication, Blainville (1826) mentioned that two specimens were sent to him by M. Florent-Prevost from the interior parts of Brazil, specifically from the province of “las Minas”. However, the original description provides no further details regarding the locality or the date of collection. A narrative emerged within the mammalian community, suggesting that the specimen was collected in the state of Minas Gerais, Brazil (e.g., Eisenberg and Redford 1999; Bonvicino et al. 2008). However, two significant challenges cast doubt on this hypothesis. Firstly, there is a geographic discrepancy, as the southern border of the state of Minas Gerais lies more than 700 km away from the nearest species of *Ctenomys* found in southern Brazil. Additionally, all other ctenomyids exhibit a contiguous distribution across South America (Maestri and Patterson 2021).

Fernandes et al. (2012) proposed an alternative hypothesis regarding the collection site of the holotype. Drawing upon the morphological similarities between *C. brasiliensis* and *C. pearsoni*, although a comprehensive comparison with other closely related species was not conducted, and considering that Uruguay, where most populations of *C. pearsoni* are found, was part of Brazil around 1826, they suggested that the city of “Minas” in Uruguay was the likely collection site of *C. brasiliensis/C. pearsoni* (Fig. 2B). Moreover, (Darwin 1839) identified a tuco-tuco as *Ctenomys brasiliensis* in Maldonado, Uruguay. This hypothesis gained significant acceptance within the community due to its greater plausibility compared to the “Minas Gerais” hypothesis (e.g., Bidau 2015; Tomasco et al. 2022).

However, upon recent examination of the type material, we discovered a label written on the wooden base of the block specimen’s frontal surface. Contrary to the information provided in the original description, this label indicates the collection locality as “Saint Paul, Brésil”, translated as São Paulo, Brazil (Fig. 1B). Surprisingly, this crucial detail appears to have been overlooked or disregarded in the previous follow-up studies. From a historical geographic standpoint, this revelation could significantly alter our understanding of the true type locality of *C. brasiliensis* and warrants further investigation. Furthermore, a DNA analysis of the type specimen has yet to be conducted due to the technical challenges associated with extracting DNA from a holotype dating back approximately 200 years. Despite these obstacles, we have chosen to pursue such analyses.

**Figure 1.**
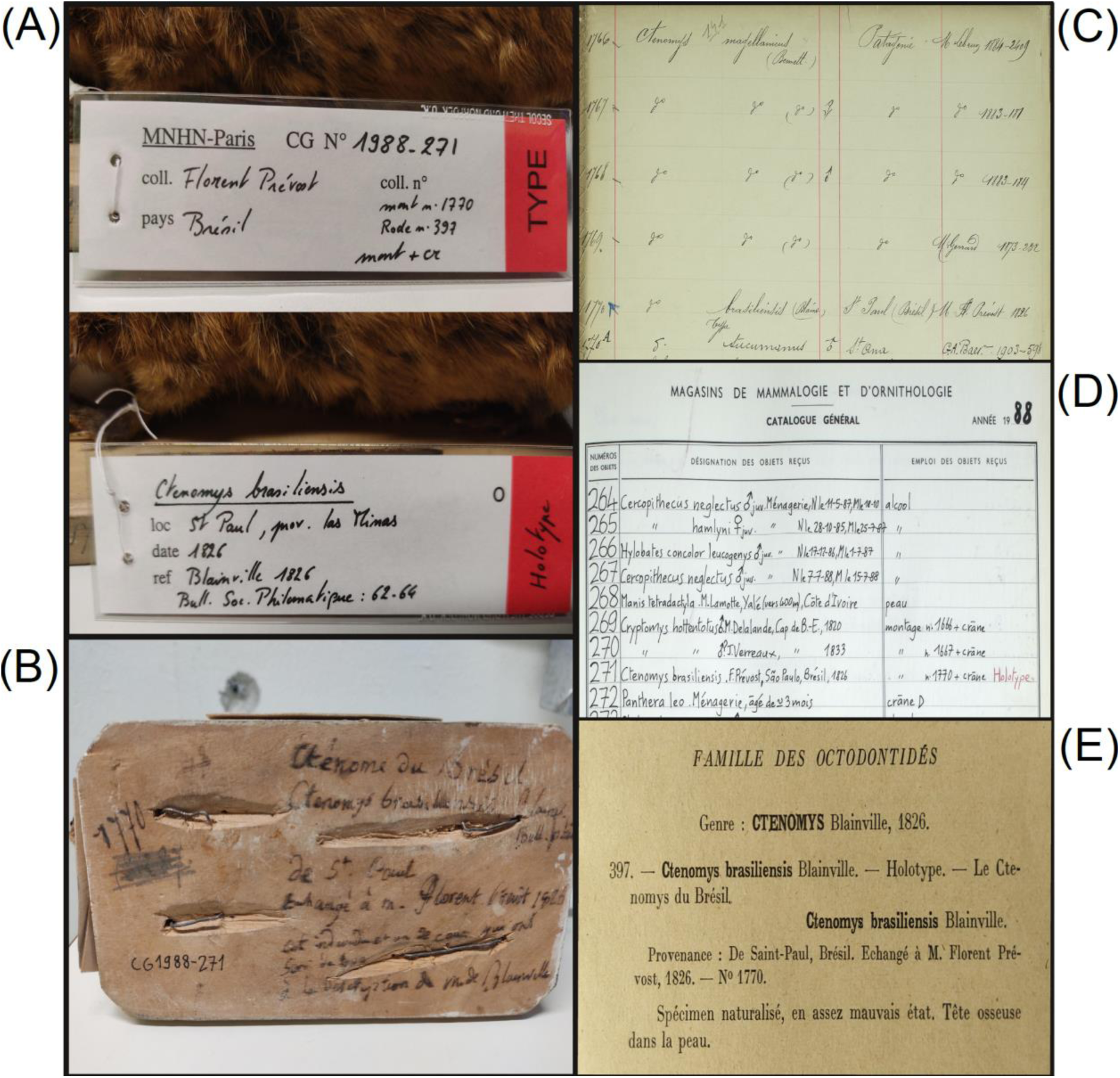
Type specimen current labels (A), and historic label (B), with information from the museum. In (C), the historical register (“Catalogue des Montages”) of the specimen in the museum. In (D), the new register from 1988, when an MNHN number was attributed to the specimen. In (E), the Rode catalog of MNHN type specimens. Photos by Violaine NC. In (B), on the wooden base, it is written “Ctenome du Brésil. Ctenomys brasiliensis. De St Paul. Echangé à Mr Florent Prevost 1826. Cet individu est un de ceux qui ont servi de type à la description de Blainville” which can be translated as “Ctenome of Brazil. Ctenomys brasiliensis. From São Paulo. Exchanged to Mr Florent Prevost 1826. This individual is one of those who served for the type description of Blainville”.

In this study, we initially sequenced the entire mitogenome of the holotype using the remaining degraded tissue available on the skull at the MNHN, employing a methodology tailored for ancient DNA. Subsequently, we conducted a comprehensive reevaluation of its cranial morphology at the geometric morphometric level, comparing it to closely related species. We then contextualized our findings within the backdrop of historical political changes in Brazil, exploring genetic affinities between the holotype and other *Ctenomys* species alongside historical information to propose a new hypothesis regarding the collection site of the holotype and its taxonomic identity. Finally, from a taxonomic perspective, we validated the type species and proposed a new synonym within the genus.

## METHODS

### DNA extraction and sequencing

The sampling of tissue from the holotype for DNA extraction was approved by a specific committee composed of curators, scientists, and conservation managers after the presentation of a scientific project and request for the museum, which was archived under the MHNH Id Colhelper 168437. Dry tissue was obtained from the cranium of the specimen. The DNA was extracted from tissue samples by the innuPREP Forensic Kit (Analytik Jena, Germany) and eluted to 55 ul of 10mM TRIS solution. The quality of extracted DNA was checked on the Agilent 2100 Bioanalyzer by the High Sensitivity DNA kit (Agilent, US). Because DNA was already fragmented to 50-200 bp fragments, we did not use DNA restriction in the next step. The genomic libraries were prepared as two independent ssDNA libraries using the xGen™ ssDNA & Low-Input DNA Library Preparation Kit (IDT, Inc., US). The libraries were checked on the Agilent 2100 Bioanalyzer by the High Sensitivity DNA kit, pooled equimolarly and sequenced by the Illumina technology in Novogene, UK (in average 5 million of 150 bp PE reads per library).

The obtained Illumina reads were mapped to the reference mitogenome of *Ctenomys leucodon* and independently to *Ctenomys rionegrensis* (GenBank accession number NC_020659 and HM544130) in Geneious 9.0.5 (Biomatters, Ltd.). The two dsDNA libraries produced 1.3 and 0,3 million PE reads in total and 3,5 and 0,8 thousand of them were mapped to the reference (with mean coverage 16 and 5,2, respectively). The consensus sequences from two different libraries were checked by eye for mismatches (they occurred only in D-loop region that was not used in follow-up analyses) and merged into a final mitogenome sequence. Alternatively, all reads from both libraries were mapped on the reference and this produced the same consensus sequence.

In addition to the holotype, specimens from six other *Ctenomys* species were included in this study (Appendix I). These species were collected in the field in Brazil and Argentina between 2005 and 2010. Genomic DNA was extracted from muscle tissue using the DNeasy Blood and Tissue Kit (Qiagen, Valencia, CA, USA) according to the manufacturer’s instructions. The quality and quantity of the extracted DNA were evaluated using a NanoDrop 2000 spectrophotometer (Thermo Fisher Scientific, Waltham, MA, USA) and a PicoGreen double-stranded DNA quantification assay kit (Life Technologies, Carlsbad, CA, USA) at the Genetics Department of the Federal University of Rio Grande do Sul. For library preparation, 250 ng of genomic DNA was sent to the Genotoul core facilities in Toulouse, France, for sequencing using the Illumina TruSeq Nano DNA Sample Prep Kit (Illumina Inc., San Diego, CA, USA). DNA fragments were sheared by ultrasonication with a Covaris M220 (Covaris Inc., Woburn, MA, USA), A-tailed, and ligated to indexed adapters. Fragments of approximately 450 bp were selected using Agencourt Ampure XP beads (Beckman Coulter, Inc.) and enriched through 8 cycles of PCR. The library was then quantified, validated, and multiplexed before being hybridized on a HiSeq 2000 flow cell using the Illumina TruSeq PE Cluster Kit v.3. Paired-end reads of 100 nucleotides were obtained using the Illumina TruSeq SBS Kit v.3 (200 cycles). Sequence quality filtering was performed using the CASAVA pipeline. Bioinformatic analysis was carried out on the Genotoul bioinformatics platform (Toulouse, France). The resulting sequences were annotated using the MITOS web server (Bernt et al. 2013) with the vertebrate mitochondrial genetic code (NCBI code Table 2).

All the mitogenome sequences generated in this study were uploaded to GenBank, with the accession numbers shown in Appendix 1.

### Phylogenetic analysis

Phylogenetic reconstruction employed the mitochondrial genome of *Ctenomys brasiliensis* (15,867 base pairs), alongside related complete mitochondrial genomes of six species of ctenomids analyzed for the first time in this study (Appendix I). To encompass a broader matrix of congeners and cover all nine species groups within the genus (Parada et al. 2011; Tomasco et al. 2024), particularly the *torquatus* group, we compiled a second dataset by gathering sequences of the Cytochrome b gene (Cyt b) from representative *Ctenomys* species (this gene is the one with the better reference dataset) obtained from GenBank (Appendix I). Since a previous BLAST search in GenBank showed greater similarity between *C. brasiliensis* and *C. minutus*, we included a third dataset with all available Cyt b sequences from *C. minutus* (Appendix I) in our analysis to explore the genealogy and topological relationships between haplotypes. Sequences were trimmed to uniform lengths and visually inspected for stop codons and reading frame shifts. Alignment was conducted using Clustal X (Thompson et al. 1997), with default alignment parameters. Sequence divergence between *C. brasiliensis* and other ctenomids was estimated using *p* distances using MEGA X (Kumar et al. 2018) for both mitogenomes and cyt b sequences.

The placement of *C. brasiliensis* within the genus was assessed through both maximum likelihood (ML) and Bayesian (BA) analyses, employing the optimal molecular evolution model (GTR + I + G) determined by the Bayesian Information Criterion (BIC) via jModeltest2 (Darriba et al. 2012). ML analysis was executed using RAxML-NG (v.0.8.1; Kozlov et al. 2019) with default parameters and the GTR substitution model (estimated by RAxML-NG), augmented by 100 bootstrap replicates to ascertain node support values. BA was performed in MrBayes 3.1 (Ronquist and Huelsenbeck 2003), involving two runs with three heated and one cold Markov chains each. Each run extended over 10 million generations, with trees sampled every 1000 generations. The initial 25% of trees were discarded as burn-in, and the remaining trees were used to generate a 50% majority-rule consensus tree with posterior probability (PP) estimates for each clade. The resulting topologies from both ML and BA analyses were visualized and edited using FigTree 1.4.4 (http://tree.bio.ed.ac.uk/software/figtree/).

The mapping of the likely collection locality of *C. brasiliensis* (Fig. 2) was done through a mapping of the range surrounding the Cyt-b haplotypes of *C. minutus* most closely related to the holotype based on a median-joining network constructed in Network 10.2 (https://www.fluxus-engineering.com).

**Figure 2.**
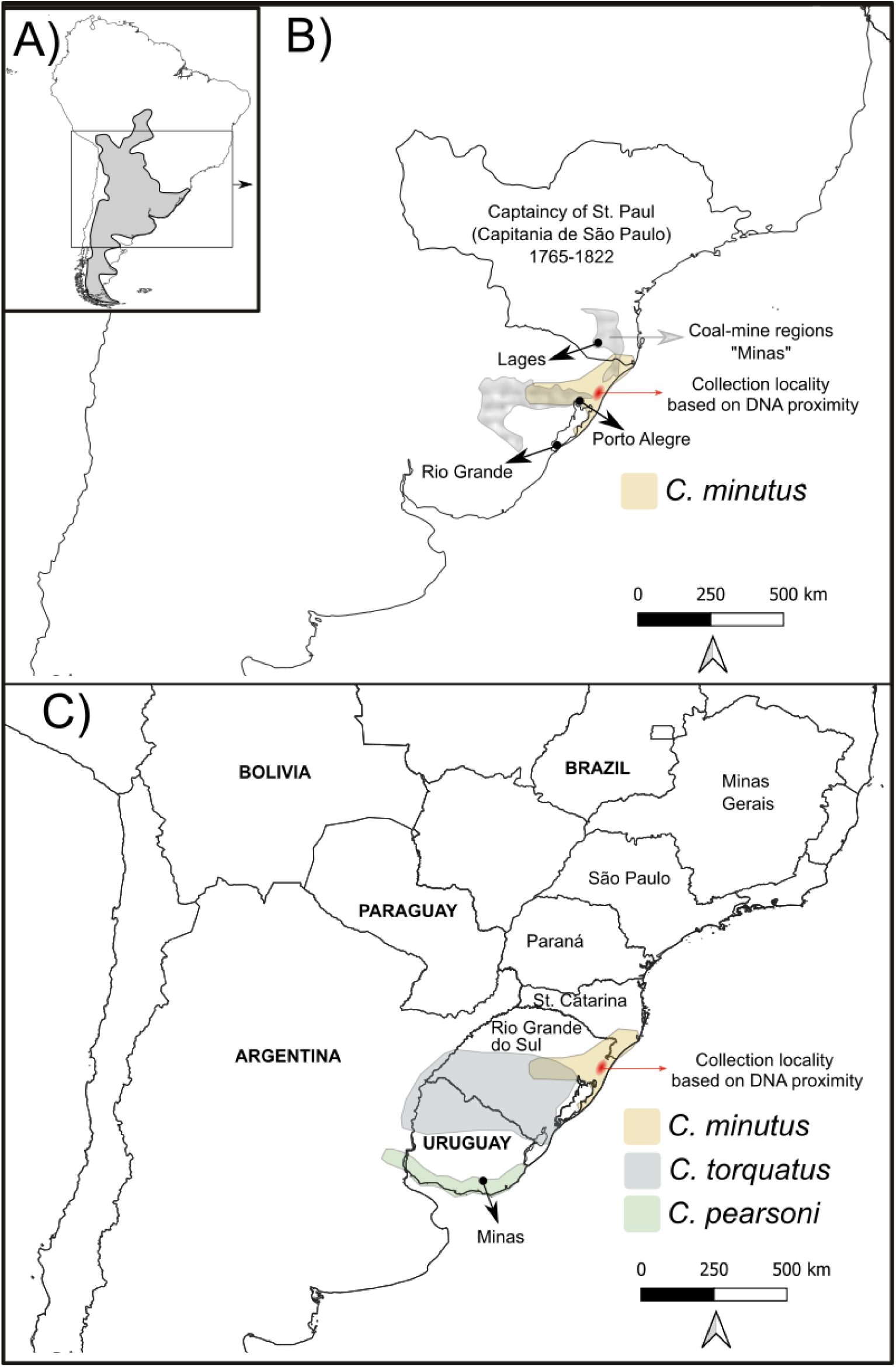
Maps. (A) Inception: map of South America with a convex polygon, in gray, showing the approximate distribution of all the species of *Ctenomys* in South America; the focal region of the next maps is within the rectangle. (B) Map showing the St. Paul captaincy from 1765 to 1822 (based on Bueno 2009). (C) The current map shows the political limits between countries and the focal Brazilian states nowadays. Coal-mine regions are delimitated according to Bunse (1984) and appear in mixed gray in panel (B); note the two “minas” regions. Range maps of current species follow Bidau (2015). The center of the red range points to the municipality of Osório-RS, the collection site of the specimen most close to the *C. brasiliensis* holotype (see also Fig. 6).

### Quantitative morphological comparison

The skull of the holotype was compared to the skulls of 165 specimens from eight species (Table 2) that form the ‘*torquatus*’ group of species (De Santi et al. 2021; D’Elía et al. 2021) using quantitative analyses based on geometric morphometrics. A list of these specimens and the museums where they are housed is provided in Supplementary Data SD3. Two-dimensional images of the crania in dorsal, lateral, and ventral views were obtained with a digital camera, at a standardized position, including a scale label. Images of the holotype were taken at the National Museum of Natural History of Paris, France (MNHN). The images of the other 165 specimens were taken from specimens deposited in eight scientific collections across the Americas (Fornel et al. 2021). Landmarks were digitized on the cranium following Fernandes et al. (2012): 19 landmarks in the dorsal view, 16 landmarks in the ventral view, and 13 landmarks in the lateral view. A description of landmark positioning and geometric morphometrics procedures is available in Supplementary Data SD3. The matrices of landmark coordinates were superimposed with a Generalized Procrustes Analysis (GPA) that removes undesirable effects of scale, position, and orientation. The dorsal and ventral views were constrained to symmetry to eliminate potential noise caused by bilateral asymmetry. Principal Component Analyses were conducted on the resulting shape matrices to explore shape variation among specimens. Procrustes distances between species were calculated for each view of the skull, considering the complete morphological information. Lastly, we averaged the Procrustes distances among views of the skull to obtain a concatenated distance from each species to *C. brasiliensis*. Landmarks were digitized using the TPS series of software (Rohlf 2015), and the statistical analyses were conducted in MorphoJ (Klingenberg 2011), and the R Software (R Core Team 2024) with the package ‘geomorph’ (Baken et al. 2021; Adams et al. 2023).

## RESULTS

### DNA results

Next-generation sequencing uncovered the mitochondrial genome of the *C. brasiliensis* holotype, which spans 15,867 base pairs and comprises 13 protein-coding genes, 22 tRNAs and 2 rRNAs. The mitochondrial genomes of six additional species were also sequenced, with lengths ranging from 15,900 to 16,400 base pairs, and displayed similar annotations. Phylogenetic analyses based on both mitogenome and Cyt b gene sequences recovered a strongly supported monophyletic group of *Ctenomys* (BS = 100; PP = 1; Fig. 3 and 4), with well-resolved relationships among the species groups generally consistent with prior studies (Parada et al. 2011).

**Figure 3.**
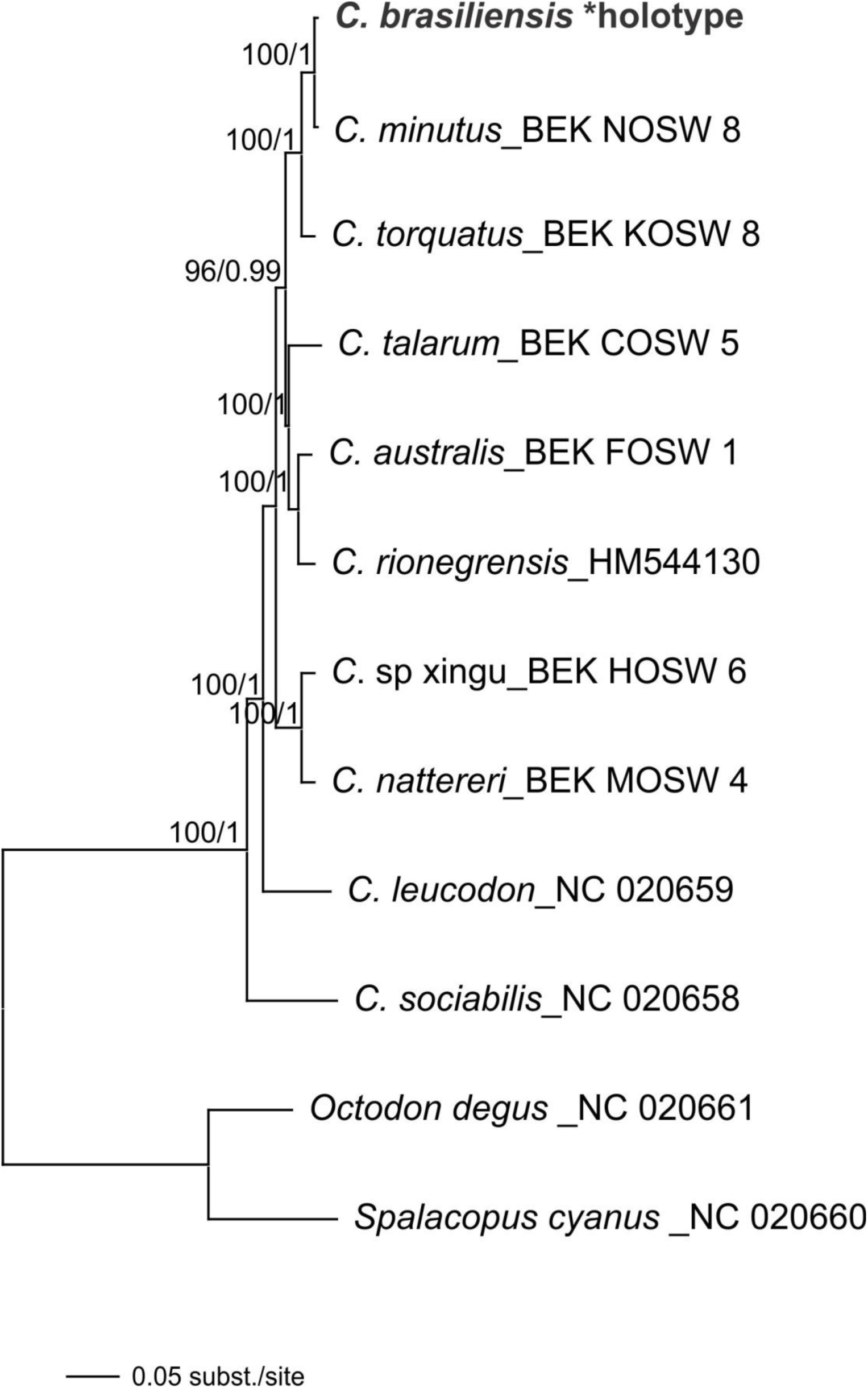
Phylogenetic analysis of mitogenomes using Maximum Likelihood. The tree was built from 10 species of *Ctenomys*, incorporating the holotype of *C. brasiliensis*, with bootstrap support values (on the left of the crossbar) and posterior probability values (on the right of the crossbar from Bayesian analysis) provided for adjacent nodes. The tree was rooted with representatives of Octodontids, *Spalacopus cyanus* and *Octodon degus*. Terminal labels indicate species names and their corresponding identification (including GenBank accession numbers).

**Figure 4.**
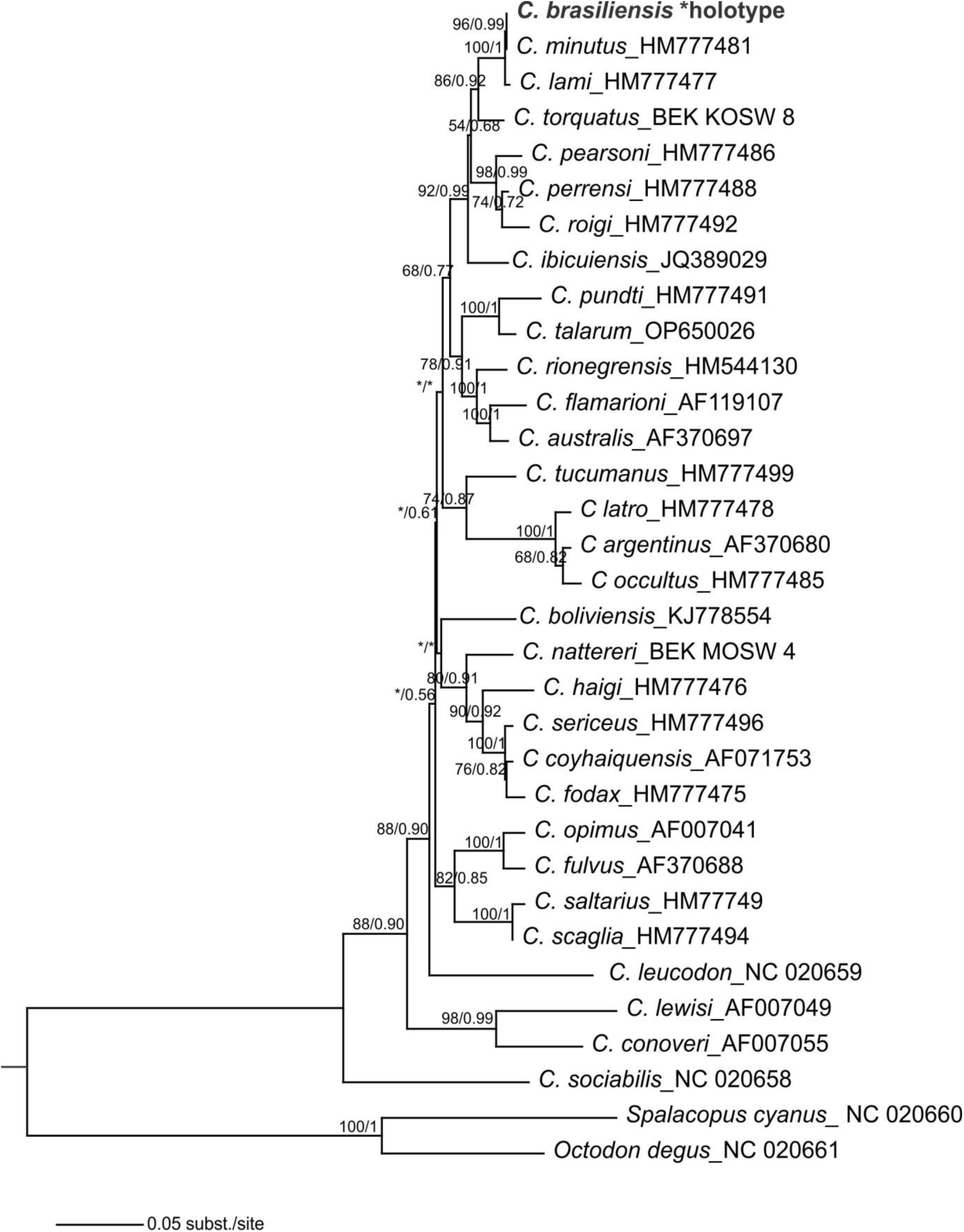
Phylogenetic analysis of mitochondrial DNA sequences using Maximum Likelihood. The tree was constructed from 31 cytochrome b gene sequences of *Ctenomys*, including the holotype of *C. brasiliensis* and representatives of all eight species groups (sensu Parada et al. 2011), with bootstrap support values (on the left of the crossbar) and posterior probability values (on the right of the crossbar from Bayesian analysis) indicated for adjacent nodes. Nodes with less than 50% posterior probability are marked with an asterisk. The tree was rooted with representatives of Octodontids, *Spalacopus cyanus*, and *Octodon degus*. Terminal labels specify species names and their corresponding GenBank accession numbers.

The *C. brasiliensis* holotype was classified within the *torquatus* group, which comprise *C. pearsoni, C. perrensi, C. roigi, C. dorbignyi, C. torquatus, C. lami, C. minutus,* and *C. ibicuiensis*, showing a close relationship to *C. minutus* in both the mitogenome tree (BS = 100, PP = 1; Fig. 3) and the Cyt b gene tree (BS = 0.96, PP = 1; Fig. 4 and Supplementary Data SD2 and SD3). Initially, the genetic distance between the holotype and *C. minutus* was minimal (almost zero), gradually increasing when compared to other species within the *torquatus* group, reaching 4% compared to *C. ibicuiensis*, the most distant species from the group, and peaking at 12% for *C. sociabilis*, the most distantly related species of *Ctenomys* (Fig. 5) for Cyt b sequences. A pairwise analysis of ctenomids using whole mitochondrial genomes yielded results consistent with those from Cyt b (Table 1). The comparative analysis with sequences of *C. minutus* showed intraspecific variation ranging from 0.2% to 2.7% (Supplementary Data SD1). The voucher CML431 was found to be 100% identical to the *C. brasiliensis* holotype, sharing the haplotype H1 in the network analysis (Fig. 6). The voucher CML431 was collected in the municipality of Osório, RS, Brazil, located at the center of the red range in Fig. 2.

**Figure 5.**
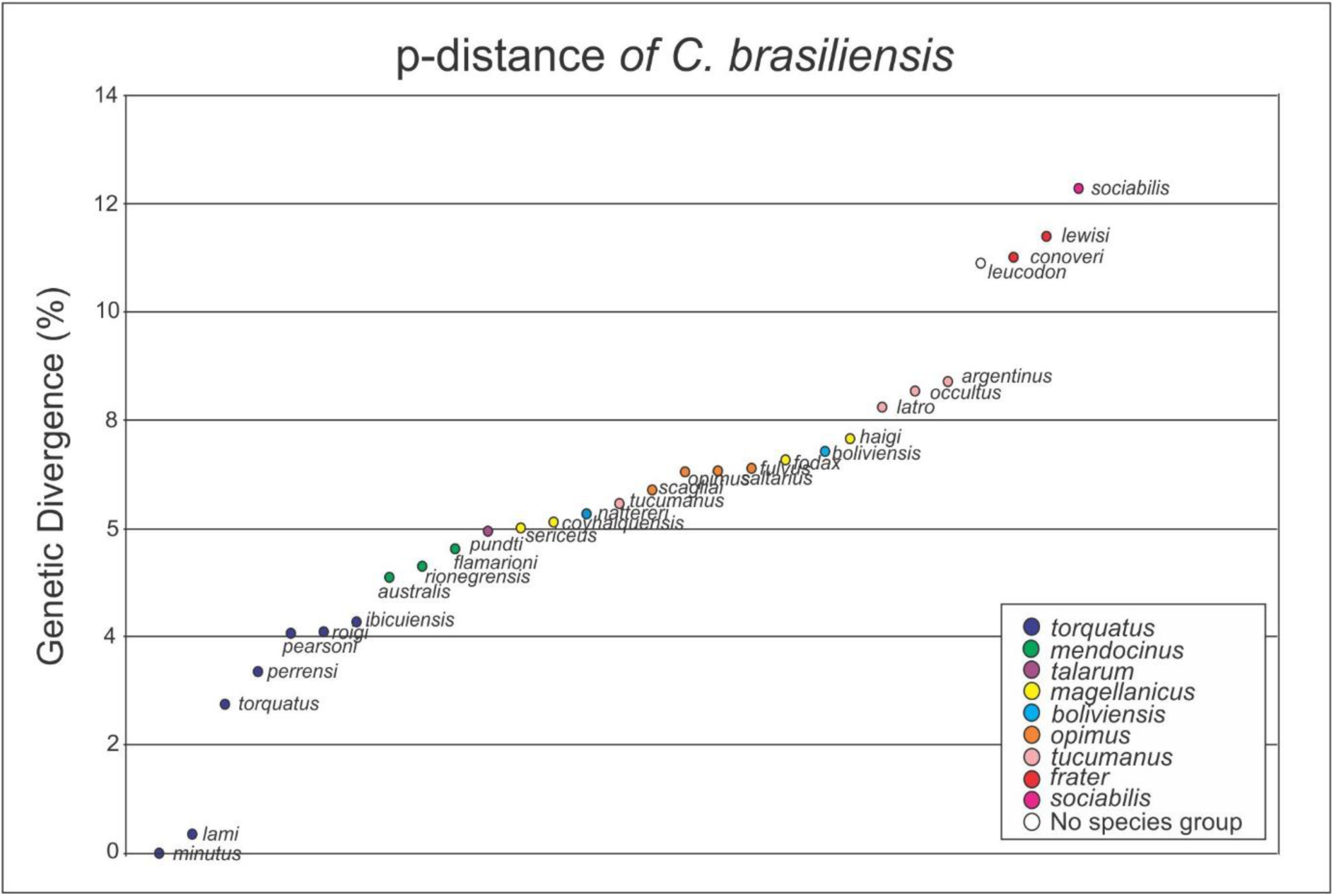
Pairwise genetic distance (%) based on cyt b sequences between *C. brasiliensis* and representatives of the nine species group (as defined by Parada et al., 2011 and Tomasco et al. 2024), indicated by colors, and ranked from the lowest to the highest values.

**Figure 6.**
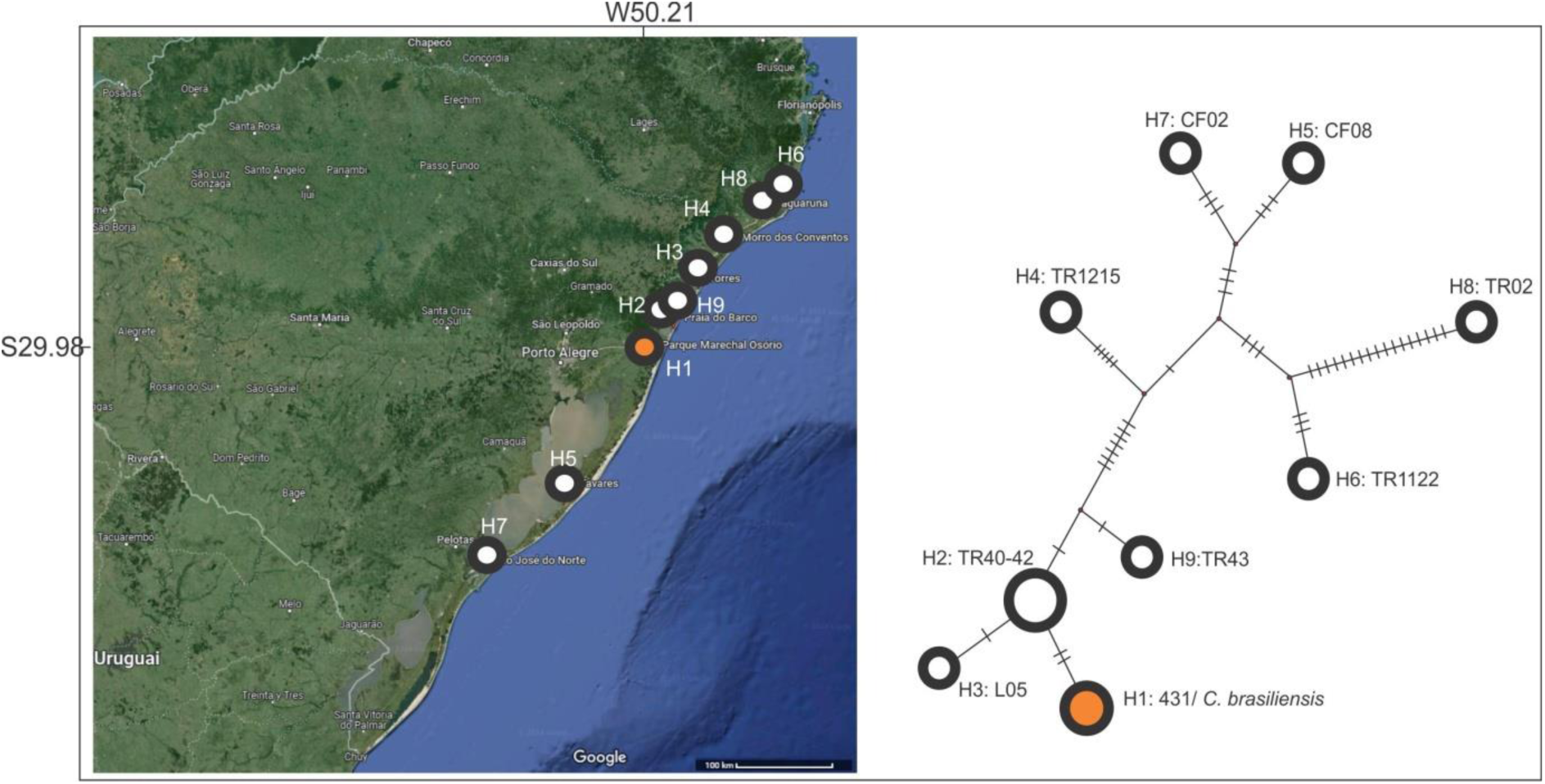
Median-joining haplotype network topology obtained with Cytochome b mtDNA data, from 11 sequences of *C. minutus* and the holotype of *C. brasiliensis*.

**Table 1.** Genetic divergence (%) of mitochondrial genomes among *Ctenomys* species. Below the diagonal are the mitogenome distances, and above the diagonal are the Cyt b distances. Values in bold indicate the pairwise divergence of *C. brasiliensis* from its congeners.

| *Ctenomys* |  |  |  |  |  |  |  |  |  |  |
| --- | --- | --- | --- | --- | --- | --- | --- | --- | --- | --- |
|  | *brasiliensis* | *minutus* | *sp. xingu* | *nattereri* | *australis* | *leucodon* | *rionegrensis* | *sociabilis* | *talarum* | *torquatus* |
| *brasiliensis* | - | **0.0** | **7.8** | **6.3** | **5.1** | **10.9** | **5.3** | **12.3** | **5.9** | **2.8** |
| *minutus* | **0.7** | - | 7.8 | 6.3 | 5.1 | 10.9 | 5.3 | 12.2 | 6.0 | 2.8 |
| *sp. xingu* | **6.4** | 6.6 | - | 2.3 | 9.1 | 12.3 | 9.8 | 14.5 | 9.7 | 8.3 |
| *nattereri* | **6.3** | 6.6 | 2.1 | - | 6.3 | 10.5 | 6.1 | 12.9 | 6.7 | 6.6 |
| *australis* | **4.3** | 4.6 | 5.9 | 5.8 | - | 11.7 | 3.4 | 13.5 | 5.0 | 5.3 |
| *leucodon* | **8.9** | 9.4 | 9.4 | 9.3 | 8.3 | - | 10.5 | 16.5 | 11.5 | 10.1 |
| *rionegrensis* | **4.6** | 5.0 | 6.1 | 6.2 | 2.5 | 8.7 | - | 14.2 | 5.1 | 5.5 |
| *sociabilis* | **11.0** | 11.5 | 11.1 | 11.1 | 10.3 | 11.9 | 10.5 | - | 13.8 | 13.0 |
| *talarum* | **5.7** | 5.5 | 7.1 | 7.0 | 4.8 | 9.8 | 5.2 | 11.5 | - | 6.0 |
| *torquatus* | **2.7** | 2.6 | 6.4 | 6.4 | 4.7 | 9.7 | 5.0 | 11.7 | 5.3 | - |

### Quantitative morphological comparison

The crania of the species from the *torquatus* group broadly overlapped in the morphospace formed by the two first Principal Components (Supplementary Data SD3), preventing a clear separation among species. Given the morphological similarity between species, quantitative morphological comparisons are hardly useful for distinguishing species. However, the average Procrustes distances between species reveal an interesting result. *Ctenomys brasiliensis* is morphologically closer to *C. minutus* (Table 2) than any other *Ctenomys* species from the *torquatus* group, despite the differences being small. Certainly, *C. brasiliensis* is closer to *C. minutus* than to *C. pearsoni* (Table 2). The second species most close to *C. brasiliensis* is *C. lami*, the sister species of *C. minutus*.

**Table 2.** Morphological comparison. Number of specimens analyzed from species of the *torquatus* group, including *Ctenomys brasiliensis*, and average Procrustes distance from *C. brasiliensis* to all other species, considering the three skull views.

| Species | Number of specimens | Procrustes distances from *C. brasiliensis* |
| --- | --- | --- |
| *C. brasiliensis* | 1 | - |
| *C. dorbignyi* | 13 | 0.0841 |
| *C. ibicuiensis* | 16 | 0.0827 |
| *C. lami* | 30 | 0.0813 |
| *C. minutus* | 30 | 0.0789 |
| *C. pearsoni* | 30 | 0.0893 |
| *C. perrensi* | 9 | 0.0845 |
| *C. roigi* | 7 | 0.0791 |
| *C. torquatus* | 30 | 0.0827 |

### Taxonomy

Accordingly, we propose to reaffirm the validation of *C. brasiliensis* as the type species of the family Ctenomyidae. In accordance with the principle of priority, we also propose reclassifying *C. minutus* as a synonym thereof.

**Family** Ctenomyidae Lesson, 1842

**Genus** *Ctenomys* Blainville, 1826

***Ctenomys brasiliensis*** Blainville, 1826 Nouv. Bull. Sci. Soc. Pjilom. Paris, 1826: 62. (type species, revalidated here)

= *Ctenomys minutus* Nehring, 1887 (new synonym, here designated).

#### Holotype

National Museum of Natural History of Paris, France, MHNH number MNHN-ZM-MO-1988-271; skin, a partial skull, collector undetermined but possibly Florent Prévost (collection number 1770). The complete mitogenome generated from this specimen was deposited in GenBank under the accession number PQ153983.

#### Type locality

Most likely: Brazil, State of Rio Grande do Sul, municipality of Osório-RS, near coordinates 50°05’W to 50°20’W longitude and 29°40’S to 30°05’S latitude. Originally described as St. Paul, Brazil, on museum records, with the additional description: las Minas, from interior parts of Brazil, appearing in the original species description.

#### Distribution

*Ctenomys brasiliensis* is distributed in localities across the Brazilian states of Santa Catarina and Rio Grande do Sul, within a range polygon of approximately 40,000 square kilometers, in southern Brazil, near the Atlantic Coast.

#### Comments

Karyotype, diagnosis, morphological characteristics, habitat and natural history, and etymology follow previous accounts on Blainville (1826) and, especially, the many follow-up studies with the synonym species *C. minutus* Nehring 1887, see, for example, Bidau (2015) and Freitas (2016).

## DISCUSSION

Our findings indicate that the holotype belongs to the same species as the specimens currently named *Ctenomys minutus* (Nehring 1887), which we now designate as a junior synonym. This result is also supported by our morphometric analyses. We begin by discussing the historical aspects related to the description of both species, clarifying taxonomic and geographic debates in light of our DNA results and historical research, and then formally validate *C. brasiliensis*.

According to Nehring (1887), the three specimens of ctenomyids from Brazil, initially identified by Th. Bischofp as *Ctenomys brasiliensis* Blainville (1826), were subsequently examined by himself (Nehring 1887). He observed that their cranial size was smaller than other ctenomyids, which led him to propose a new species, *C. minutus*, supposedly named after these characteristics related to the small size. Nehring (1887, 1900) also noted that other European specialists, including Oldfield Thomas, immediately questioned the validity of his proposed new species, suggesting that these specimens were immature individuals, hence their smaller cranial dimensions. Nehring (1887) further compared the type specimens with those of other ctenomyid species, such as *Ctenomys torquatus*, recommending further investigation. In a later study, Nehring (1900) compared *C. minutus* with *C. torquatus* and other species, including *Ctenomys pundti*, which he newly proposed. Surprisingly, he did not include *C. brasiliensis* in this comparative analysis, and subsequent studies debating its validity based on body size have remained inconclusive. Some researchers have suggested synonymizing *C. brasiliensis* with *C. torquatus* (e.g. Wagner 1848; Vieira 1955; Voss 1973), *C. minutus* (e.g. Reig et al. 1965) or *C. pearsoni* Lessa and Langguth 1983 (e.g. Fernandes et al. 2012). Because of its poor original characterization by Blainville (1826), limited subsequent material confined to type specimens, and imprecise collection locality in Brazil with unknown collector and date, *C. brasiliensis* has been regarded by some as *species inquirendae* (e.g. (Cichchino et al. 2000). Notably, even geometric morphometric analysis involving cranial measurements across several landmarks has shown minimal, if any, differences among these (e.g. Fernandes et al. 2012). In our study, which included the type specimen of *C. brasiliensis* for comparison, no clear differences were found in size and shape. Therefore, taxonomic decisions regarding this issue cannot rely solely on morphometry, whether through linear measurements or geometric morphometric analyses.

The identification of the type locality of *C. brasiliensis* may have been subject to misinterpretation over time. The reference to “las Minas”, the sole information on the specimens’ collection location provided by Blainville (1826), which has been associated with “Minas Gerais state” by some authors, appears unfounded. No ctenomid specimens have been collected along the Brazilian coast north of the municipality of Jaguaruna in present-day Santa Catarina, the northern limit of the current *C. minutus* range (Fig. 2C). Additionally, the collection locality labeled as “St. Paul = São Paulo”, on the oldest label written on the type specimens’ wooden block (Fig. 1) does not appear in Blainville’s original description in 1826. At that time, the St. Paul Captaincy—“Capitania de São Paulo” (Bueno 2009)—comprised a significant portion of the Brazilian east coast, but did not extend beyond the northern boundary of present-day Minas Gerais (Fig. 2). Moreover, the captaincy of St. Paul extended south as far as the towns of Lages and Laguna. Therefore, its geographic scope did not encompass the southernmost region under Portuguese influence, which was the “Provincia do Rio Grande de São Pedro” (current Rio Grande do Sul State and Uruguay). Consequently, attributing the type locality to “Minas” in Lavajeda, the southernmost part of Uruguay (Fernandes et al. 2012), is also considered implausible.

Furthermore, the southern extent of the captaincy of St. Paul from 1765 to 1822 does not overlap with the geographical range of *C. torquatus* or *C. pearsoni*. This is not the case for *C. minutus*, which inhabits the coastal region up to the Laguna area, overlapping with the southernmost region of the former captaincy of St. Paul (Fig. 2). The type locality of *C. minutus*, described as “Mundo Novo” (= New World, at the time of its description by Nehring (1887)) encompass a large, vaguely defined area south of the Serra do Mar mountains in Rio Grande do Sul, which is located northeast to Porto Alegre city and roughly coincides in part with present distribution of *C. minutus* (Fig. 1). This region was initially viewed by Portuguese authorities as promising for economic development, colonization, and agricultural exploration. Significant colonization of this area began after 1826, culminating in its elevation to the status of “Freguesia de Taquara do Mundo Novo,” comprising at least six present-day municipalities (Taquara, Riozinho, Três Coroas, Igrejinha, Rolante, and Paranhana at a later date) (Reinheimer 2005). Interestingly, Nehring (1887) speculated a more specific type locality for *C. minutus* near the Tramandaí River, originating in the northeastern part of the aforementioned region. This particular area coincides with the closest DNA sequence found for *C. minutus* (voucher 431) compared to *C. brasiliensis* in our current study (marked in red in Fig. 2), sharing the Cyt b-haplotype H1. Voucher 431 has a diploid number of 2n=46a. *Ctenomys minutus* exhibits high karyotype polymorphism, with chromosomal variation categorized into eight parental karyotypes (2n=50a, 48c, 46a, 48a, 42, 46b, 48b, and 50b (Freitas 1997; Freygang et al. 2004). According to Lopes et al. (2013), the 2n=46a mtDNA haplogroup belongs to the Coastal and Barros Lake regions, including specimens collected at Gaivota Beach, Passo de Torres, Guarita Beach, Barco Beach, and Tramandaí.

It is worth noting that the current geographical distribution of *C. minutus* partially overlaps with historical low-altitude coal mines discovered in southern Brazil (Fig. 2). These mines are known since the end of the XVIII century (Witkowski 2016) in Rio Grande do Sul (RS) state, with coal extraction dating back at least to 1853 (Dahne 1893; Da Silva 2007), and are still active in municipalities like Arroio dos Ratos, Butiá (formerly Minas do Butiá) and Minas do Leão in RS, as well as those of Tubarão and Criciúma in Santa Catarina (SC) state. They are intersected by two main ancient trails used for land travel between São Paulo and Rio Grande de São Pedro provinces, leading to Viamão city: traversing the highlands via Lages and Taquara, and the other along the lowlands passing through present-day Laguna (SC) and Torres municipalities (RS) (for a detailed description, see Bueno 2009; Beier and Cintra 2016). The latter route, for instance, was traveled by August Saint-Hillaire, a naturalist from the MNHN, during his exploratory journey to the region in 1822 (Saint-Hilaire 1887). Thus, it cannot be ruled out that the term “las Minas”, considered the type locality in the original description of *C. brasiliensis*, might alternatively refer to these specific coal mines rather than those discussed earlier.

Finally, the successful retrieval of *C. brasiliensis* mitogenome sequences in this study, which is genetically almost identical to *C. minutus*, and regarding Cyt b only, it nested entirely within the sequences of *C. minutus* (Supplementary Data S2) constitutes a significant discovery that strongly supports the arguments presented above. The species most closely related to *C. minutus* is *C. lami*, with a 0.3% of genetic difference, both nested within the same clade of *C. brasiliensis*. However, *C. minutus* and *C. lami* belong to an evolutionary complex and form a hybrid zone (Lopes et al. 2013; Freitas et al. 2021). Our analysis relies on a single sequence of *C. lami* from GenBank, which is questionably identified due to the difficulty in distinguishing these species using mtDNA data. Consequently, we consider it necessary to obtain more data with reliable identification from the parental area of *C. lami* for further comparison with *C. brasiliensis* in this taxonomic context.

## ACKNOWLEDGEMENTS

We are grateful to the MNHN for allowing us to sequence the DNA and take pictures of the type specimen. We thank Fabiano A. Fernades for *C. brasiliensis* skull’s images. RM was supported by CAPES and CNPq (302990/2022-4). GRPM was supported by CNPq fellowship. RF was supported by CNPq (309745/2023-3). TROF was supported by CAPES, CNPq and FAPERGS. Thanks to all curators and collection managers that provided access to *Ctenomys* specimens: Enrique González (MUNHINA), Olga B. Vacaro and Esperança A. Varela (MACN), A. Damián Romero (MMP), James L. Patton, Eileen A. Lacey, and Christopher Conroy (MVZ), Eileen Westwig (AMNH), and Bruce D. Patterson (FMNH).

## Supplementary Data

Supplementary Data SD1.— Genetic divergence table between *C. brasiliensis* and *C.minutus*.

Supplementary Data SD2.— Cyt b tree of the *torquatus* group. Supplementary Data SD3.— Detailed morphological analyses and results.

## Appendix I

Specimens used in genetic analyses, including representatives of the nine phylogenetic species groups (sensu Parada et al. 2011 and Tomasco et al. 2024) and an outgroup, with GenBank accession numbers. Species group: ‘*opimus*’: *C. fulvus*, AF370688; *C. opimus*, AF007041; *C. saltarius*, HM777493; *C. scagliai*, HM777494. ‘*mendocinus*’: *C. flamarioni*, AF119107; *C. rionegrensis*, HM544130/ AF119106; *C. australis*, AF370697/ BEK_FOSW_1. ‘*talarum*’: *C. pundti*, HM777491; *C. talarum*, TBD/BEK_COSW_5, OP650026. ‘*torquatus*’: *C. brasiliensis*, PQ153983; *C. pearsoni*, HM777486; *C. perrensis*, HM777488; *C. minutus* TBD/ BEK_NOSW_8, HM777481 (CML 431), HM777482 (TR40), JQ389046 (cf02), JQ389047 (cf08), JQ389048 (L05), JQ389049 (TR1122), JQ389050 (TR1215), HM777482 (TR40), MK452114 (TR41), MK452115 (TR42), MK452116 (TR43); *C. lami*, HM777477; *C. torquatus* TBD/ BEK_KOSW_8, JQ389045; *C. roigi*, HM777492; *C. ibicuiensis*, JQ389029. ‘*magellanicus*’: *C. sericeus*, HM777496; *C. haigi*, HM777476; *C. coyhaiquensis*, AF071753; *C. fodax*, HM777475. ‘*tucumanus*’: *C. latro*, HM777478; *C. occultus*, HM777485; *C. argentinus*, AF370680; *C. tucumanus*, HM777499. ‘*boliviensis*’: *C. nattereri*, TDB/ BEK_MOSW_4, MK895145; *C. boliviensis*, KJ778554; *C. sp. ‘xingu’*, TBD/ BEK_HOSW_6, MK895247. ‘*frater*’: *C. lewisi*, AF007049; *C. conoveri*, AF007055. ‘*sociabilis*’: *C. sociabilis*, NC020658. ‘Incertae’: *C. leucodon*, NC020659. Outgroup: *Octodon degus*, NC020661; *Spalacopus cyanus*, NC020660.

## REFERENCES

de Abreu-Jr EF, Pavan SE, Tsuchiya MTN, Wilson DE, Percequillo AR, Maldonado JE. 2020. Museomics of tree squirrels: a dense taxon sampling of mitogenomes reveals hidden diversity, phenotypic convergence, and the need of a taxonomic overhaul. BMC Evol Biol. 20(1):77. doi:10.1186/s12862-020-01639-y.

Adams DC, Collyer ML, Kaliontzopoulou A, Baken EK. 2023. Geomorph: Software for geometric morphometric analyses. R package version 4.0.6. <https://cran.r-project.org/package=geomorph>.

Alvarado-Larios R, Teta P, Cuello P, Jayat JP, Tarquino-Carbonell AP, D’Elía G, Cornejo P, Ojeda AA. 2024. A new living species of the genus Ctenomys (Rodentia: Ctenomyidae) from central-western Argentina. Vertebr Zool. 74:193–207. doi:10.3897/vz.74.e115242.

Baken EK, Collyer ML, Kaliontzopoulou A, Adams DC. 2021. geomorph v4.0 and gmShiny: Enhanced analytics and a new graphical interface for a comprehensive morphometric experience. Methods Ecol Evol. 12(12):2355–2363. doi:10.1111/2041-210X.13723.

Beier JR, Cintra JP. 2016. O MAPA DA CAPITANIA DE SÃO PAULO DE WILHELM LUDWIG VON ESCHWEGE: UMA ANÁLISE CARTOGRÁFICA.

Bidau CJ. 2015. Family Ctenomyidae. In: Mammals of South America, Vol. 2: Rodents. The University of Chicago Press, Chicago,: Patton JL, Pardiñas UFJ, D’Elía G (eds).

Blainville HMD. 1826. Sur une nouvelle espèce de Rongeur fouisseur du Brésil. Bull Soc Philomath Paris.:62–64.

Bonvicino CR, de Oliveira JA, D’Andrea PS. 2008. Guia dos roedores do Brasil, com chaves para gêneros baseadas em caracteres externos. Rio Jan Cent Pan-Am Febre Aft - OPASOMS.:120.

Brook F, Tomasco IH, González B, Martin GM. 2022. A New Species of Ctenomys (Rodentia: Ctenomyidae) from Patagonia Related to C. sociabilis. J Mamm Evol. 29(1):237–258. doi:10.1007/s10914-021-09570-9.

Bueno BPS. 2009. Dilatação dos confins: caminhos, vilas e cidades na formação da Capitania de São Paulo (1532-1822). An Mus Paul História E Cult Mater. 17(2):251–294. doi:10.1590/S0101-47142009000200013.

Castañeda-Rico S, Edwards CW, Hawkins MTR, Maldonado JE. 2022. Museomics and the holotype of a critically endangered cricetid rodent provide key evidence of an undescribed genus. Front Ecol Evol. 10:930356. doi:10.3389/fevo.2022.930356.

Cichchino AC, Castro DC, Baldo JC. 2000. Elenco de los Phthiraptera (Insecta) hallados em distintas poblaciones locales de Ctenomys (Rodentia: Octodontidae) de Argentina, Uruguay, Paraguay, Bolivia y Brasil. Papéis Avulsos Zool.:197–211.

Da Silva CE. 2007. Nas profundezas da Terra: um estudo sobre a região carbonífera do Rio Grande do Sul (1883-1945).

Dahne ES. 1893. A mineração de carvão e as concessões da companhia no Estado do Rio Grande do Sul. Cia Estrada Ferro E Minas São Jeronymo Estabel Typogr Gundlanch.

Darriba D, Taboada GL, Doallo R, Posada D. 2012. jModelTest 2: more models, new heuristics and parallel computing. Nat Methods. 9(8):772–772. doi:10.1038/nmeth.2109.

Darwin C. 1839. Narrative of the surveying voyages of His Majesty’s Ships Adventure and Beagle between the years 1826 and 1836, describing their examination of the southern shores of South America, and the Beagle’s circumnavigation of the globe. London: Henry Colburn: Journal and Remarks.

De Santi NA, Verzi DH, Olivares AI, Piñero P, Álvarez A, Morgan CC. 2021. A new Pleistocene *Ctenomys* and divergence dating of the hyperdiverse South American rodent family Ctenomyidae. J Syst Palaeontol. 19(5):377–392. doi:10.1080/14772019.2021.1910583.

D’Elía G, Teta P, Lessa EP. 2021. A Short Overview of the Systematics of Ctenomys: Species Limits and Phylogenetic Relationships. In: Tuco-tucos: an evolutionary approach to the diversity of a Neotropical subterranean rodent. Springer Nature Switzerland: Freitas, T.R.O., Gonçalves, G.L., Maestri, R. 10.1007/978-3-030-61679-3.

Eisenberg JF, Redford KH. 1999. Mammals of the neotropics, the central neotropics. Chicago: The University of Chicago Press.

Fernandes FA, Fornel R, Freitas TRO. 2012. Ctenomys brasiliensis Blainville (Rodentia: Ctenomyidae): clarifying the geographic placement of the type species of the genus Ctenomys. Zootaxa. 3272(1). doi:10.11646/zootaxa.3272.1.4. [accessed 2024 Jun 6]. https://mapress.com/zt/article/view/zootaxa.3272.1.4.

Fornel R, Maestri R, Cordeiro-Estrela P, De Freitas TRO. 2021. Skull Shape and Size Diversification in the Genus Ctenomys (Rodentia: Ctenomyidae). In: Tuco-tucos: an evolutionary approach to the diversity of a Neotropical subterranean rodent. Springer Nature Switzerland: Freitas, T.R.O., Gonçalves, G.L., Maestri, R. 10.1007/978-3-030-61679-3.

Freitas TRO. 1997. Chromosome polymorphism in Ctenomys minutus (Rodentia– Octodontidae). Rev Bras Genética.:1–7.

Freitas TROD, Gonçalves GL, Maestri R, editors. 2021. Tuco-Tucos: An Evolutionary Approach to the Diversity of a Neotropical Subterranean Rodent. Cham: Springer International Publishing. [accessed 2024 Jun 6]. https://link.springer.com/10.1007/978-3-030-61679-3.

Freygang CC, Marinho JR, Freitas TRO. 2004. New karyotypes and some considerations about the chromosomal diversication of. Ctenomys minutus (Rodentia: Ctenomyidae) on the coastal plain of the Brazilian state of Rio Grande do Sul. Genética.:125–132.

Klingenberg CP. 2011. MorphoJ: an integrated software package for geometric morphometrics. Mol Ecol Resour. 11(2):353–357. doi:10.1111/j.1755-0998.2010.02924.x.

Kozlov AM, Darriba D, Flouri T, Morel B, Stamatakis A. 2019. RAxML-NG: a fast, scalable and user-friendly tool for maximum likelihood phylogenetic inference. Bioinformatics. 35(21):4453–4455. doi:10.1093/bioinformatics/btz305.

Kumar S, Stecher G, Li M, Knyaz C, Tamura K. 2018. MEGA X: Molecular Evolutionary Genetics Analysis across Computing Platforms. Mol Biol Evol. 35(6):1547–1549. doi:10.1093/molbev/msy096.

Lessa EP, Langguth A. 1983. Ctenomys pearsoni, n. sp. (Rodentia: Octodontidae), del Uruguay. Resúmenes Comuniciones Las Jorn Cienc Nat Urug.:86–88.

Lesson RP. 1842. Nouveau Tableau du Règne Animal. Mammifères. Paris: Arthus-Bertrand.

Lopes CM, Ximenes SSF, Gava A, de Freitas TRO. 2013. The role of chromosomal rearrangements and geographical barriers in the divergence of lineages in a South American subterranean rodent (Rodentia: Ctenomyidae: Ctenomys minutus). Heredity. 111(4):293–305. doi:10.1038/hdy.2013.49.

Maestri R, Patterson BD. 2021. Geographical and macroecological patterns of tuco-tucos. In: Tuco-tucos: an evolutionary approach to the diversity of a Neotropical subterranean rodent. Springer Nature Switzerland: Freitas, T.R.O., Gonçalves, G.L., Maestri, R. 10.1007/978-3-030-61679-3.

Mammal Diversity Database. 2024. Mammal Diversity Database (v1.12.1) [Data set]. 10.5281/zenodo.10595931.

Mapelli FJ, Teta P, Contreras F, Pereyra D, Priotto JW, Coda JA. 2023. Looking under stones: A new Ctenomys species from the rocky foothills of the Sierras Grandes of central Argentina. J Mamm Evol. 30(1):281–298. doi:10.1007/s10914-022-09634-4.

McGuire JA, Cotoras DD, O’Connell B, Lawalata SZS, Wang-Claypool CY, Stubbs A, Huang X, Wogan GOU, Hykin SM, Reilly SB, et al. 2018. Squeezing water from a stone: high-throughput sequencing from a 145-year old holotype resolves (barely) a cryptic species problem in flying lizards. PeerJ. 6:e4470. doi:10.7717/peerj.4470.

Nations JA, Giarla TC, Morni MA, William Dee J, Swanson MT, Hiller AE, Khan FAA, Esselstyn JA. 2022. Molecular data from the holotype of the enigmatic Bornean Black Shrew, Suncus ater Medway, 1965 (Soricidae, Crocidurinae), place it in the genus Palawanosorex. ZooKeys. 1137:17–31. doi:10.3897/zookeys.1137.94217.

Nehring A. 1887. Über eine Ctenomys-Art aus Rio Grande do Sul (Süd Brasilien). Sitzungsb Ges Naturf Fr.:45–57.

Nehring A. 1900. Über die Schädel von Ctenomys minutusNhrg., Ct. torquatus Licht und Ct. pundti Nhrg. SitzungsbGes Naturf Fr.:201–210.

Pagès M, Chaval Y, Herbreteau V, Waengsothorn S, Cosson J-F, Hugot P, Morand S, Michaux J. 2010. Revisiting the taxonomy of the Rattini tribe: a phylogeny-based delimitation of species boundaries.

Parada A, D’Elía G, Bidau CJ, Lessa EP. 2011. Species groups and the evolutionary diversification of tuco-tucos, genus *Ctenomys* (Rodentia: Ctenomyidae). J Mammal. 92(3):671–682. doi:10.1644/10-MAMM-A-121.1.

R Core Team. 2024. R Core Team. https://www.Rproject.org/.

Reig OA, Contreras JR, Piantanida MJ. 1965. Contribución a la elucidación sistemática de las entidades del género Ctenomys (Rodentia, Octodontidae). I. Relaciones de parentesco entre muestras de ocho poblaciones de tuco-tucos inferidas del estudio estadístico de variables del fenotipo y su correlación con las características del cariotipo. Contrb Cient Ser Zool.:299–352.

Reinheimer D. 2005. Terra, gente e fé: Aspectos históricos de Taquara do Mundo Novo. Taquara, Faccat.

Rohlf F. 2015. The tps series of software. Hystrix Ital J Mammal. 26(1). doi:10.4404/hystrix-26.1-11264. [accessed 2024 Jun 6]. 10.4404/hystrix-26.1-11264.

Ronquist F, Huelsenbeck JP. 2003. MrBayes 3: Bayesian phylogenetic inference under mixed models. Bioinformatics. 19(12):1572–1574. doi:10.1093/bioinformatics/btg180.

Ruelas D, Pacheco V. 2022. New morphological data and phylogenetic position of the rare short-tailed opossum Monodelphis ronaldi (Didelphimorphia: Didelphidae) with new records. Mamm Biol. 102(1):189–204. doi:10.1007/s42991-021-00219-x.

Saint-Hilaire A. 1887. Voyage à Rio-Grande do Sul (Brésil). Orléans, H. Herluison.

Sánchez RT, Tomasco IH, Díaz MM, Barquez RM. 2023. Review of three neglected species of Ctenomys (Rodentia: Ctenomyidae) from Argentina. J Mammal. 104(3):578–590. doi:10.1093/jmammal/gyad001.

Tammone MN. 2024. A new species of *Ctenomys* (Rodentia, Ctenomyidae) from the pre-Andean regions of Mendoza Province, Argentina. Rowe K, editor. J Mammal. 105(3):609– 620. doi:10.1093/jmammal/gyae024.

Teta P, Jayat JP, Alvarado-Larios R, Ojeda AA, Cuello P, D’Elía G. 2023. An appraisal of the species richness of the Ctenomys mendocinus species group (Rodentia: Ctenomyidae), with the description of two new species from the Andean slopes of west-central Argentina. Vertebr Zool. 73:451–474. doi:10.3897/vz.73.e101065.

Thompson JD, Gibson TJ, Plewniak F, Jeanmougin F, Higgins DG. 1997. The CLUSTAL_X windows interface: Flexible strategies for multiple sequence alignment aided by quality analysis tools. :4876–4882.

Tomasco IH, Ceballos SG, Austrich A, Brook F, Caraballo DA, Fernández GP, Lanzone C, Mora MS, Parada A, Sánchez RT, et al. 2024. Underground speciation: Unraveling the systematics and evolution of the highly diverse tuco-tucos (genus *Ctenomys*) with genomic data. Mol Phylogenet Evol. 199:108163. doi:10.1016/j.ympev.2024.108163.

Tomasco IH, Giorello FM, Boullosa N, Feijoo M, Lanzone C, Lessa EP. 2022. The contribution of incomplete lineage sorting and introgression to the evolutionary history of the fast-evolving genus Ctenomys (Rodentia, Ctenomyidae). Mol Phylogenet Evol. 176:107593. doi:10.1016/j.ympev.2022.107593.

Upham NS, Patterson BD. 2015. Evolution of caviomorph rodents: a complete phylogeny and timetree for living genera. In: Biology of caviomorph rodents: diversity and evolution. Buenos Aires: SAREM Series A: Vassallo AI, Antenucci D.

Verzi DH, De Santi NA, Olivares AI, Morgan CC, Álvarez A. 2021. The History of Ctenomys in the Fossil Record: A Young Radiation of an Ancient Family. In: Freitas TRO de, Gonçalves GL, Maestri R, editors. Tuco-Tucos: An Evolutionary Approach to the Diversity of a Neotropical Subterranean Rodent. Cham: Springer International Publishing. p. 3–15. [accessed 2024 Jul 19]. 10.1007/978-3-030-61679-3_1.

Verzi DH, De Santi NA, Olivares AI, Morgan CC, Basso NG, Brook F. 2023. A new species of the highly polytypic South American rodent Ctenomys increases the diversity of the magellanicus clade. Vertebr Zool. 73:289–312. doi:10.3897/vz.73.e96656.

Vieira CC. 1955. Lista Remissiva dos Mamíferos do Brasil. Arq Zool Est São Paulo.:341– 374.

Voss WA. 1973. Ensaio de lista sistemática dos mamíferos do Rio Grande do Sul. Pesquisas.:1–35.

Wagner R. 1848. Beiträge zur Kennteniss der Arten von Ctenomys. Arch F Naturg.:72–78.

Witkowski A. 2016. HOSPITAL SARMENTO LEITE: PATRIMÔNIO CULTURAL DA REGIÃO CARBONÍFERA DO BAIXO JACUÍ – RIO GRANDE DO SUL. In: A História e o campo do patrimônio: desafios e perspectivas. Porto Alegre RS: T Nacional História e Patrimônio Cultural ANPUH Brasil. (I seminário nacional de história e patrimônio cultural). p. 193–207.

Yuan SC, Malekos E, Cuellar-Gempeler C, Hawkins MTR. 2022. Population genetic analysis of the Humboldt’s flying squirrel using high-throughput sequencing. J Mammal. 103(2):287–302. doi:10.1093/jmammal/gyac002.

