## Supplementary Data for "Ancient DNA Clarifies the Identity and Geographic Origin of the holotype of the genus *Ctenomys*"

**Supplementary Data SD1**

**Table SD1.** Comparative genetic divergence (p-distance) based on Cyt b sequences from *C. minutus* and *C. brasiliensis* (holotype). Genbank accession numbers and voucher of *C. minutus* are indicated.

|  | Holotype | HM777481  CML431 | MK452114  TR41 | MK452115  TR42 | JQ389048  L05 | MK452116  TR43 | HM777482  TR40 | JQ389050  TR1215 | JQ389047  cf08 | JQ389049  TR1122 | JQ389046  cf02 | HM777483  TR02 |
| --- | --- | --- | --- | --- | --- | --- | --- | --- | --- | --- | --- | --- |
| Holotype |  |  |  |  |  |  |  |  |  |  |  |  |
| HM777481_CML431 | 0.000 |  |  |  |  |  |  |  |  |  |  |  |
| MK452114_TR41 | 0.002 | 0.002 |  |  |  |  |  |  |  |  |  |  |
| MK452115_TR42 | 0.002 | 0.002 | 0.000 |  |  |  |  |  |  |  |  |  |
| JQ389048_L05 | 0.003 | 0.003 | 0.001 | 0.001 |  |  |  |  |  |  |  |  |
| MK452116_TR43 | 0.004 | 0.004 | 0.002 | 0.002 | 0.003 |  |  |  |  |  |  |  |
| HM777482_TR40 | 0.004 | 0.004 | 0.000 | 0.000 | 0.001 | 0.002 |  |  |  |  |  |  |
| JQ389050_TR1215 | 0.014 | 0.014 | 0.013 | 0.013 | 0.014 | 0.013 | 0.013 |  |  |  |  |  |
| JQ389047_cf08 | 0.016 | 0.016 | 0.014 | 0.014 | 0.015 | 0.014 | 0.014 | 0.013 |  |  |  |  |
| JQ389049_TR1122 | 0.018 | 0.018 | 0.016 | 0.016 | 0.017 | 0.016 | 0.016 | 0.013 | 0.013 |  |  |  |
| JQ389046_cf02 | 0.019 | 0.019 | 0.016 | 0.016 | 0.018 | 0.016 | 0.017 | 0.014 | 0.006 | 0.014 |  |  |
| HM777483_TR02 | 0.027 | 0.027 | 0.028 | 0.028 | 0.027 | 0.028 | 0.027 | 0.023 | 0.026 | 0.019 | 0.027 |  |

**Supplementary Data SD2**

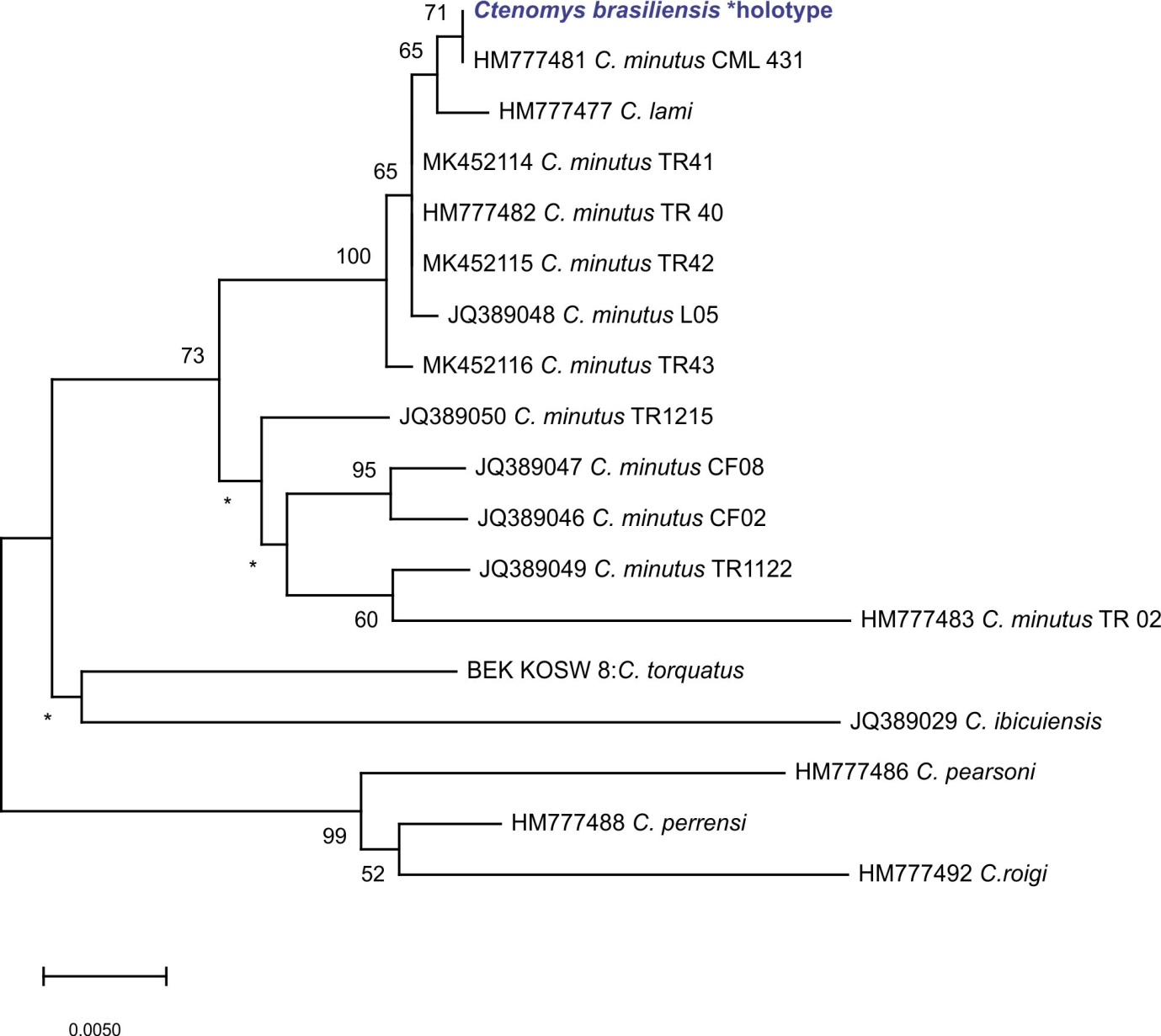

**Figure SD2.** Phylogenetic analysis of mitochondrial DNA sequences using the Maximum Likelihood method. The resulting tree was constructed from the Cytochrome b gene sequences of *Ctenomys brasiliensis*, all available sequences from *C. minutus,* and representatives of the *torquatus* species group (sensu Parada et al. 2011). Bootstrap support values are indicated at the corresponding nodes, with nodes showing less than 50% posterior probability marked with an asterisk. Terminal labels include species names and their respective GenBank accession numbers.

**Supplementary Data SD3**

**Detailed morphological procedures and results**

**Data**

The 165 specimens of *Ctenomys* compared to *Ctenomys brasiliensis*, along with their scientific collection/museum number, are listed below. Museum labels/identifiers below correspond to:

TR = Departamento de Genética, Universidade Federal do Rio Grande do Sul, Porto Alegre, Brazil

AMNH = American Museum of Natural History, New York, USA

FMNH = Field Museum of Natural History, Chicago, USA

MACN = Museo Argentino de Ciencias Naturales “Bernardino Rivadavia”, Buenos Aires, Argentina

MUNHINA = Museo Nacional de Historia Natural y Antropología, Montevideo, Uruguay

MMPMa = Museo Municipal de Ciencias Naturales “Lorenzo Scaglia”, Mar del Plata, Argentina

MVZ = Museum of Vertebrate Zoology, University of California, Berkeley, USA

| **Identifier in the scientific collection** | **Species** |
| --- | --- |
| TR-77 | *C.lami* |
| TR-78 | *C.lami* |
| TR-79 | *C.lami* |
| TR-86 | *C.lami* |
| TR-87 | *C.lami* |
| TR-88 | *C.lami* |
| TR-89 | *C.lami* |
| TR-95 | *C.lami* |
| TR-97 | *C.lami* |
| TR-98 | *C.lami* |
| TR-1070 | *C.lami* |
| TR-109 | *C.lami* |
| TR-110 | *C.lami* |
| TR-111 | *C.lami* |
| TR-112 | *C.lami* |
| TR-113 | *C.lami* |
| TR-114 | *C.lami* |
| TR-115 | *C.lami* |
| TR-131 | *C.lami* |
| TR-132 | *C.lami* |
| TR-133 | *C.lami* |
| TR-134 | *C.lami* |
| TR-340 | *C.lami* |
| TR-344 | *C.lami* |
| TR-346 | *C.lami* |
| TR-349 | *C.lami* |
| TR-353 | *C.lami* |
| TR-356 | *C.lami* |
| TR-357 | *C.lami* |
| TR-360 | *C.lami* |
| TR-49 | *C.minutus* |
| TR-50 | *C.minutus* |
| TR-90 | *C.minutus* |
| TR-106 | *C.minutus* |
| TR-123 | *C.minutus* |
| TR-124 | *C.minutus* |
| TR-126 | *C.minutus* |
| TR-127 | *C.minutus* |
| TR-128 | *C.minutus* |
| TR-130 | *C.minutus* |
| TR-229 | *C.minutus* |
| TR-230 | *C.minutus* |
| TR-231 | *C.minutus* |
| TR-234 | *C.minutus* |
| TR-281 | *C.minutus* |
| TR-288 | *C.minutus* |
| TR-289 | *C.minutus* |
| TR-290 | *C.minutus* |
| TR-291 | *C.minutus* |
| TR-293 | *C.minutus* |
| TR-294 | *C.minutus* |
| TR-299 | *C.minutus* |
| TR-300 | *C.minutus* |
| TR-302 | *C.minutus* |
| TR-304 | *C.minutus* |
| TR-312 | *C.minutus* |
| TR-314 | *C.minutus* |
| TR-324 | *C.minutus* |
| TR-326 | *C.minutus* |
| TR-327 | *C.minutus* |
| TR-69 | *C.torquatus* |
| TR-70 | *C.torquatus* |
| TR-71 | *C.torquatus* |
| TR-92 | *C.torquatus* |
| TR-93 | *C.torquatus* |
| TR-96 | *C.torquatus* |
| TR-116 | *C.torquatus* |
| TR-117 | *C.torquatus* |
| TR-122 | *C.torquatus* |
| TR-144 | *C.torquatus* |
| TR-148 | *C.torquatus* |
| TR-1505 | *C.torquatus* |
| TR-152 | *C.torquatus* |
| TR-153 | *C.torquatus* |
| TR-154 | *C.torquatus* |
| TR-155 | *C.torquatus* |
| TR-156 | *C.torquatus* |
| TR-2026 | *C.torquatus* |
| TR-2247 | *C.torquatus* |
| TR-2254 | *C.torquatus* |
| AMNH-206485 | *C.torquatus* |
| AMNH-206486 | *C.torquatus* |
| AMNH-206487 | *C.torquatus* |
| AMNH-206488 | *C.torquatus* |
| AMNH-206489 | *C.torquatus* |
| AMNH-206490 | *C.torquatus* |
| TR 397 | *C.torquatus* |
| TR 398 | *C.torquatus* |
| TR 955 | *C.torquatus* |
| TR 956 | *C.torquatus* |
| AMNH-206512 | *C.pearsoni* |
| AMNH-206513 | *C.pearsoni* |
| AMNH-206514 | *C.pearsoni* |
| AMNH-206515 | *C.pearsoni* |
| AMNH-206516 | *C.pearsoni* |
| AMNH-206517 | *C.pearsoni* |
| AMNH-206518 | *C.pearsoni* |
| AMNH-206519 | *C.pearsoni* |
| AMNH-206520 | *C.pearsoni* |
| AMNH-206521 | *C.pearsoni* |
| AMNH-206522 | *C.pearsoni* |
| AMNH-206523 | *C.pearsoni* |
| AMNH-206525 | *C.pearsoni* |
| AMNH-206526 | *C.pearsoni* |
| AMNH-206527 | *C.pearsoni* |
| AMNH-206529 | *C.pearsoni* |
| AMNH-206532 | *C.pearsoni* |
| AMNH-206534 | *C.pearsoni* |
| AMNH-206535 | *C.pearsoni* |
| AMNH-206536 | *C.pearsoni* |
| AMNH-206545 | *C.pearsoni* |
| AMNH-206546 | *C.pearsoni* |
| FMNH-29302 | *C.pearsoni* |
| MACN-19562 | *C.pearsoni* |
| MACN-19563 | *C.pearsoni* |
| MACN-19569 | *C.pearsoni* |
| MACN-19587 | *C.pearsoni* |
| MUNHINA-1838 | *C.pearsoni* |
| MUNHINA-1841 | *C.pearsoni* |
| MUNHINA-2249 | *C.pearsoni* |
| MMPMa-2413 | *C.perrensi* |
| MMPMa-2414 | *C.perrensi* |
| MMPMa-2437 | *C.perrensi* |
| MMPMa-2440 | *C.perrensi* |
| MMPMa-2441 | *C.perrensi* |
| MMPMa-2447 | *C.perrensi* |
| MMPMa-3418 | *C.perrensi* |
| MVZ-179151 | *C.perrensi* |
| MVZ-179152 | *C.perrensi* |
| MMPMa-3424 | *C.dorbignyi* |
| MMPMa-3425 | *C.dorbignyi* |
| MMPMa-3426 | *C.dorbignyi* |
| MMPMa-3427 | *C.dorbignyi* |
| MMPMa-3428 | *C.dorbignyi* |
| MMPMa-3429 | *C.dorbignyi* |
| MMPMa-3432 | *C.dorbignyi* |
| MMPMa-3452 | *C.dorbignyi* |
| MMPMa-3455 | *C.dorbignyi* |
| MMPMa-3456 | *C.dorbignyi* |
| MMPMa-3457 | *C.dorbignyi* |
| MMPMa-3459 | *C.dorbignyi* |
| MVZ-179149 | *C.dorbignyi* |
| MMPMa-2410 | *C.roigi* |
| MMPMa-2411 | *C.roigi* |
| MMPMa-2442 | *C.roigi* |
| MMPMa-3417 | *C.roigi* |
| MMPMa-3461 | *C.roigi* |
| MVZ-179153 | *C.roigi* |
| MVZ-179154 | *C.roigi* |
| TR-1065 | *C.ibicuiensis* |
| TR-1066 | *C.ibicuiensis* |
| TR-1067 | *C.ibicuiensis* |
| TR-1068 | *C.ibicuiensis* |
| TR-1070 | *C.ibicuiensis* |
| TR-1072 | *C.ibicuiensis* |
| TR-1073 | *C.ibicuiensis* |
| TR-1074 | *C.ibicuiensis* |
| TR-1505 | *C.ibicuiensis* |
| TR-1506 | *C.ibicuiensis* |
| TR-1507 | *C.ibicuiensis* |
| TR-1508 | *C.ibicuiensis* |
| TR-1509 | *C.ibicuiensis* |
| TR-1510 | *C.ibicuiensis* |
| TR-1513 | *C.ibicuiensis* |
| TR-1517 | *C.ibicuiensis* |

**Methods**

The position of the landmarks is shown in Figure SD3-1, below. Landmarks were defined according to Fernandes et al. (2012), as follows:

Dorsal view of the cranium: 1- anterior tip of the suture between premaxillas; 2-3- anterolateral extremity of incisor alveolus; 4- anterior extremity of the suture between nasals; 5-6- anteriormost point of the suture between nasal and premaxilla; 7-8- anteriormost point of the root of zygomatic arch; 9- suture between nasals and frontals; 10-11- anterolateral extremity of lacrimal bone; 12-13- point of least width between frontals; 14-15- tip of extremity of superior jugal process; 16-17- anterolateral extremity of suture between frontal and squamosal; 18-19- lateral extremity of suture between jugal and squamosal.

Ventral view of the cranium: 1- anterior tip of suture between premaxillas; 2-3- anterolateral extremity of incisor alveolus; 4-5- lateral edge of incisive foramen in suture between premaxilla and maxilla; 6-7- anteriormost point of root of zygomatic arch; 8-9- anteriormost point of orbit in inferior zygomatic root; 10-11- anteriormost point of premolar alveolus; 12-13- posterior extremity of III molar alveolus; 14- posterior extremity of suture between palatines; 15-16- anteriormost point of intersection between jugal and squamosal.

Lateral view of the cranium: 1- anteriormost point of premaxilla; 2- posteriormost point of incisor alveolus; 3- inferiormost point of incisor alveolus; 4- anterior tip of nasal; 5- anteriormost point of the suture between nasal and premaxilla; 6- suture between premaxilla, maxilla and frontal in superior zygomatic root; 7- inferiormost point of suture between lacrimal and maxilla; 8- inferiormost point of infraorbital foramen in inferior zygomatic root; 9- inferiormost point of suture between premaxilla and maxilla; 10- anteriormost point of premolar alveolus; 11- extremity of superior jugal process; 12- extremity of inferior jugal process; 13- extremity of posterior jugal process.

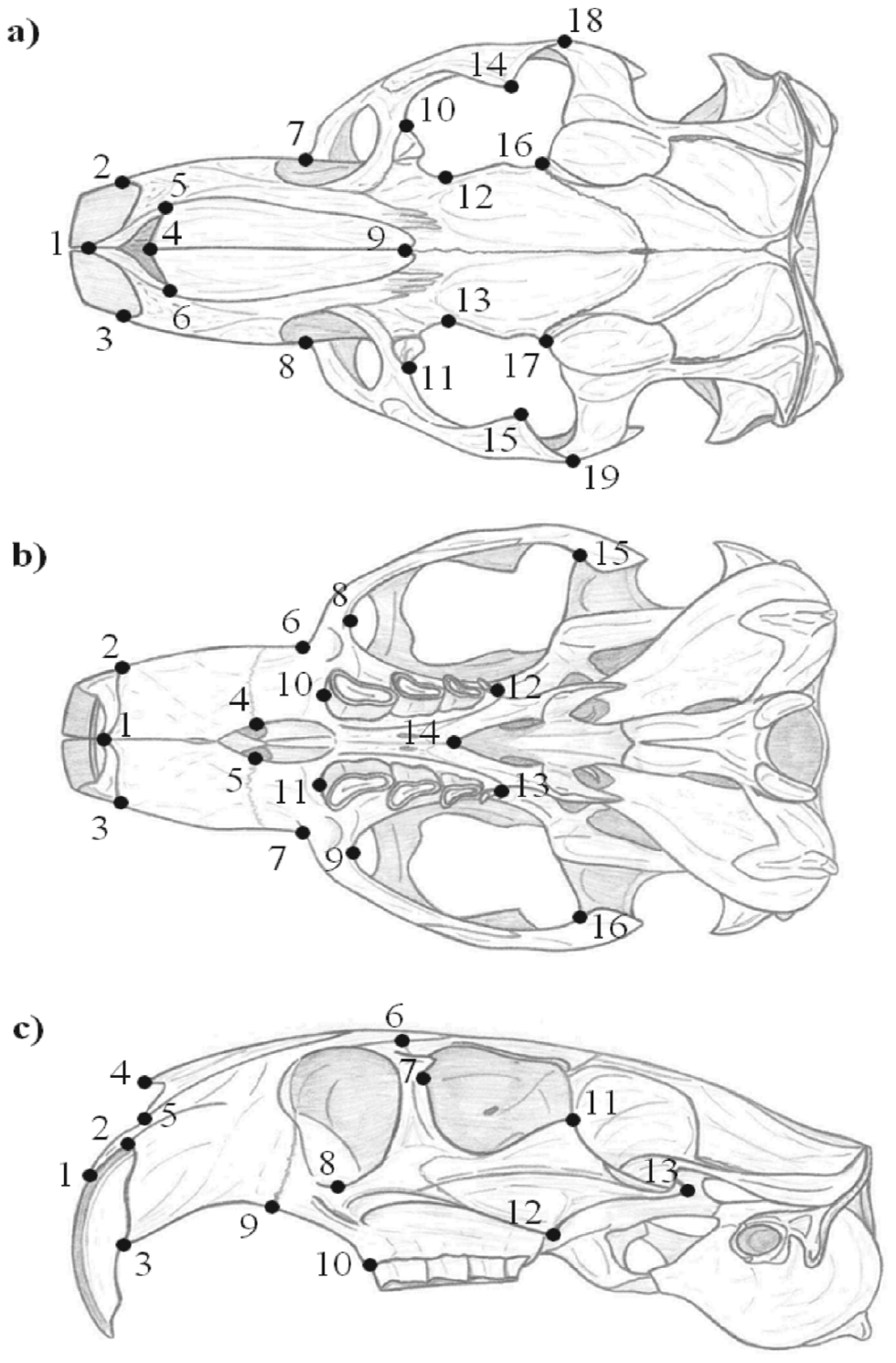

**Figure SD3-1**. Landmark positioning in the skull of the specimens in dorsal (a), ventral (b), and lateral (c) view. Drawings by Rodrigo Fornel. Modified from Fornel et al. 2010.

**Results**

Principal component analyses reveal an overlap between species of the *torquatus* group in the two first principal components, shown below. The two first components of the dorsal view explained 36,86% and 17,67% respectively; the two first components of the ventral view explained 32,40% and 16,98% respectively; and the two first components of the lateral view explained 22,91% and 15,97% respectively.

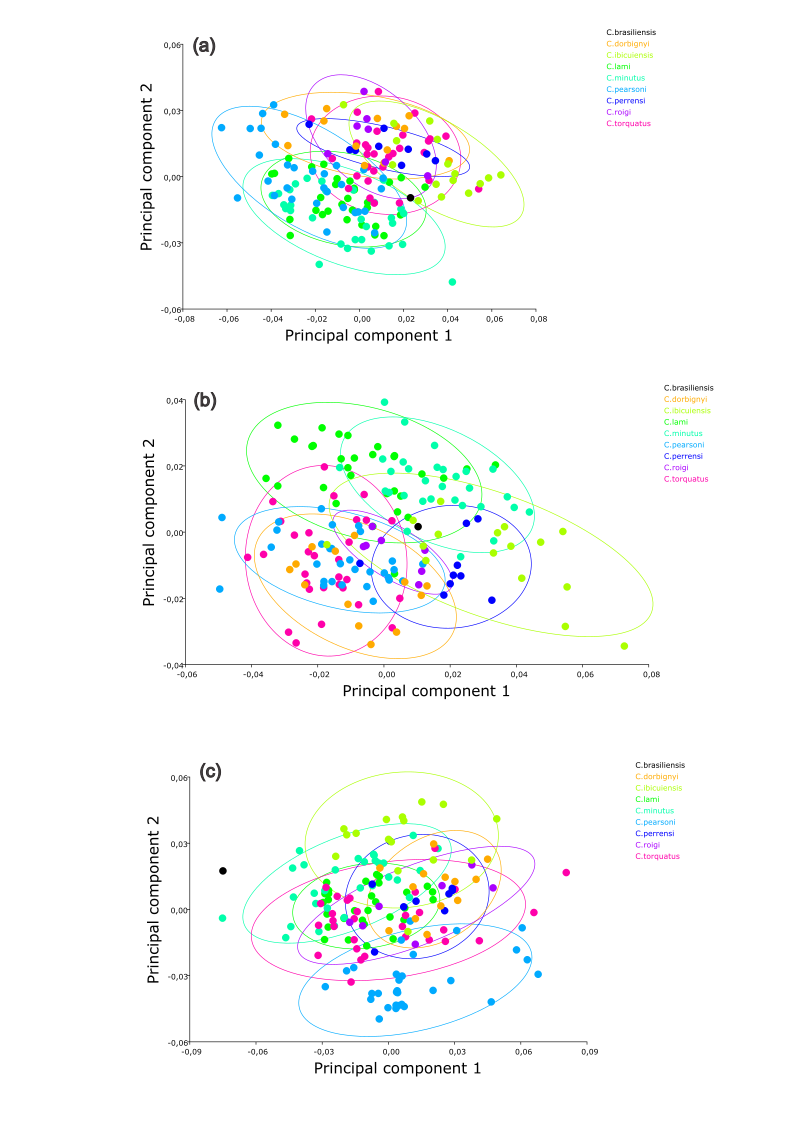

**Figure SD3-2**. Principal component analysis showing the skull shape variation among *Ctenomys* species of the *torquatus* group, including the holotype of *C. brasiliensis* (the black dot). Elipses of 90% confidence for each species are shown. In (a) the dorsal view, in (b) the ventral view, and in (c) the lateral view of the skull.
